# Tn-seq of *Thermus thermophilus* genome reveals unexpected tolerance to insertions in bacterial common essential genes

**DOI:** 10.1101/2025.04.03.647156

**Authors:** Cristina L. Gómez-Campo, Marc Gost, Bruna Fernanda Silva de Sousa, Laura Álvarez, José Berenguer, Modesto Redrejo-Rodríguez, Mario Mencía

## Abstract

A large library based on a Tn5 minitransposon carrying a thermostable kanamycin resistance gene was prepared using *Thermus thermophilus* HB27 genomic DNA as target. To increase the yield of transformants, DNA from the *in vitro* transposition reaction was amplified using isothermal multiple displacement amplification. The resulting product was first transformed into the high-transformation efficiency *ΔaddAB/ppol* strain and then into a wild-type HB27 strain. Tn-seq analysis of the libraries showed that almost all genes contained insertions and the distribution of the number of insertions per gene was unimodal, unlike the bimodal distribution reported in most Tn-seq analyses, thus hindering the discrimination of required or essential genes. Upon comparing the Tn-seq results with gene conservation in pangenomic analysis from *Thermus thermophilus* to *Deiococcota* levels, as well as with available HB27 RNA-seq data, we observed a very low correlation between core genes or gene transcription levels and Tn-seq insertion frequency. Notably, many genes largely deemed part of the essential bacterial core, supporting critical cellular pathways, showed relatively high transposon insertion numbers. In the case of DNA repair routes, which are essential but somewhat redundant, our results align well with previously published essentiality data, indicating that many genes are dispensable and permissive to insertions. The analysis of these striking results in the context of *Thermus* biology suggests that the polyploidy of the *Thermus* genome and the differential stability of proteins may explain the apparent non-essentiality of key genes.

**Importance:** The Tn-seq technique, random insertion of transposons into an organism’s genome followed by deep sequencing of the resulting population, has enabled precise identification of genes necessary for that organism to live under specified conditions. We have performed Tn-seq on the extremophilic bacterium *Thermus thermophilus* to gain information on the genes required for thriving at high temperature. Unexpectedly, we detected transposon insertions in the practical totality of the genes of this organism, hindering a clear discrimination of essential genes. Furthermore, a comparison with core genes derived from a pangenome analysis, spanning from the *Thermus* genus up to the phylum level, failed to establish a correlation between gene conservation and Tn-seq results. The polyploidy of the genome of this organism, along with potential regulatory mechanisms, may explain why commonly essential bacterial genes tolerate transposon insertions, a characteristic that appears particular for this organism.

## Introduction

Bacteria are among the most diverse and ubiquitous forms of life on Earth. They can be found in nearly every habitat, from the human gut to the deep ocean floors and even in extreme environments such as hot springs and radioactive waste. Among the extremophilic bacteria, *Thermus thermophilus* has become a cornerstone model organism for studying life at high temperature, with a growing body of knowledge about its biology and great usefulness as a source of enzymes for biotechnology (Cava, Hidalgo, and Berenguer 2009).

The genus *Thermus* is characterized by extreme thermophilic bacteria that grow at temperatures ranging from 55 to 80 °C (da Costa, Rainey, and Nobre 2006). Together with the radiation-resistant family *Deinococcaceae* and others, they form the *Deinococcus-Thermus* phylum (Deinococcota) (Ho et al. 2016). The most studied strains of *T. thermophilus* (Tth from here on) are HB27 and HB8. The HB27 genome is approximately 2.1 Mbp in size, divided between the chromosome (1.89 Mbp) and the pTT27 megaplasmid (0.23 Mbp) (Henne et al. 2004). Its predicted 2263 genes are tightly packaged within a genome that has a high GC content (69%). Another important aspect of Tth genetics is its polyploidy, with 5-8 copies of the genome present during the whole cell cycle (N. Ohtani, Tomita, and Itaya 2010).

*Thermus* species typically have aerobic heterotrophic metabolisms, and their genomes encode numerous degradative enzymes, some of which are exported to the extracellular milieu. Interestingly, *Thermus* genomes are known to be highly dynamic, containing a considerable number of insertion sequences, transposons and defense mechanisms against phages and other elements (Blesa, Averhoff, and Berenguer 2018). This is not surprising given that *Thermus* species are constitutively competent, with a very efficient machinery for environmental DNA uptake (Schwarzenlander and Averhoff 2006), which facilitates horizontal gene transfer phenomena but also ramps up the requirement for robust defense mechanisms against potentially harmful exogenous nucleic acids (Lopatina et al. 2019).

Transposon sequencing (Tn-seq) is a powerful genetic technique used to study the gene function on a genome-wide scale. This method involves the random insertion of antibiotic resistance marker-bearing transposons into bacterial DNA, followed by deep-sequencing of the surviving bacterial population to map transposon insertion sites (Van Opijnen, Bodi, and Camilli 2009). This technique allows the high-throughput identification of genes essential for bacterial survival, virulence, and/or adaptation under specific conditions (Van Opijnen and Levin 2020). Two general strategies can be followed for generating a transposon-mutagenized bacterial population. *In vivo* transposition methods are typically very efficient but require a transposition system that functions efficiently in the target species (Cho 2021). In the case of extremophilic bacteria, *in vitro* transposition is more feasible. This approach involves inserting the transposon cassette into purified genomic DNA, which is then introduced into the study strain via transformation or electroporation. Subsequently, the endogenous DNA repair machinery recombines the flanking sequences, thus incorporating the transposon at the corresponding sites in the gemone (Guschinskaya et al. 2016). In many bacterial species, the distribution of transposon insertion indices per gene typically results in a bimodal distribution: the initial sharp peak represents genes where transposon insertion is lethal, while the second, broader peak comprises genes that tolerate mutations without impairing bacterial viability under the tested conditions (Solaimanpour, Sarmiento, and Mrázek 2015).

In recent years, powered by the ever-growing number of sequences in the databases, the study of pangenomes has revealed the dynamic nature of bacterial genomes (Cummins et al. 2022). A pangenome includes the entire set of genes present in all strains of a species, comprising both the core genome, which consists on the genes shared by all strains, and the accessory genome, which contains genes present in some but not all strains. The core genome highlights the set of genes, a proportion of them essential for survival, that define the common functions of a given species, or, put in another way, what constitutes the distinctive *chassis* of the species (Tonkin-Hill, Corander, y Parkhill 2023; Tettelin et al. 2008). The accessory genome, on the other hand, is the sum of all the genes that a given species is known to bear, in any of its strains, and therefore could eventually utilize. Understanding the interplay between bacterial diversity, genetic adaptation, and environmental specialization requires an integrated approach that combines multiple scientific techniques and perspectives. Here, we explore the genetic basis of thermophily and the evolutionary strategies employed by Tth to thrive in extreme environments.

In this article we utilize the Tn5 transposon donor repeats flanking a thermostable kanamycin resistance cassette to perform *in vitro* transposition over Tth HB27 genomic DNA. To maximize the number of transformants, the transposition reaction product was amplified by isothermal multiple displacement amplification using Φ29 DNA polymerase, and the resultaning DNA was introduced onto the high transformation and high recombination capacity strain Δ*ppol/addAB* (Verdú et al. 2022) to generate a high-diversity transposition library. In the absence of a detectable bimodal curve, we used unsupervised clustering to group genes of lower, intermediate and higher insertion rate, as less permissive, intermediate, and highly permissive genes, respectively. We then compared the Tn-seq results with the gene conservation data we inferred from the pangenome of *Thermus*, *Thermaceae* and *Deinococcota* genomes available in the Refseq database. Unexpectedly, although we could detect an overall pattern of less insertions in genes coding for essential functions, like protein synthesis or DNA replication and repair, many genes known to be essential across studied bacteria and conserved at the family or class taxonomic level had a high hit rate in our study, and, in general, we found very few genes with total absence of transposons. We propose that the high tolerance to Tn insertions within expected essential genes could be related with the polyploidy of Tth, which would allow the coexistence of disrupted and functional alleles. These results provide a new perspective on the use of Tn- seq to determine gene essentiality in bacteria.

## Results

### Library construction and Tn-seq

In this study we developed a procedure for selecting random insertion mutants of Tth based on a gene cassette encoding a thermostable resistance to kanamycin flanked by the Tn5 transposase recognition sites (ME sites, Table S1) (Kia et al. 2017). The transposition of this cassette was performed onto Tth HB27 genomic DNA using a modified Tn5 transposase. To increase the concentration of DNA to be used for transformation, the reaction product of the transposition was amplified using an isothermal multiple displacement reaction kit based on the highly processive Φ29 DNA polymerase. To further maximize the number of transformants, the amplified DNA was initially transformed into a strain (Δ*ppol/addAB*) that lacks the genes coding for the primase-polymerase (Ppol, TTC0656) and the AddAB helicase-nuclease (TTC0638-0639), along with additional mutations (Verdú et al. 2022). This strain is genetically stable and shows a 100-fold increase in transformation efficiency for integrative constructs compared to the wild type (García-Quintans et al. 2020). This aproach enabled the generation of a library of 1,4x10^5^ kanamycin-resistant clones (Ppol_1 library). To regenerate the library in the wild-type HB27 strain, genomic DNA was prepared from the Ppol_1 library and transformed onto the wild-type strain, resulting in the HB27_1 library, with of 7.2x10^5^ clones. Both libraries were plated from the stocks to get the replicates Ppol_2 (3x10^5^ colonies) and HB27_2 (1.8 x10^5^ colonies). The libraries were not subjected to competitive outgrowth selection in liquid culture, as the constitutive natural competence of Tth, combined with its recombinational repair efficiency, would likely lead to a loss of population diversity.

Genomic DNA extracted from pooled libraries was processed for Tn-seq (see Materials and Methods) and sequencing was performed using an Illumina MiSeq system with 150 cycles. Between 1.5 and 3.2 million reads were obtained, depending on the library (Table 1, Table S2), of which between 40 to 90% passed the quality and duplication filters. Mapping the filtered and processed reads to the HB27 reference genome showed that the initial parental library (Ppol_1) had a somewhat lower alignment rate than the other three libraries, but all the libraries had similar genome coverage depth and breadth (Figure S1). Accordingly, the distribution of genome coverage showed that transposons were inserted all across the genome but with peaks of higher frequency of insertion also dispersed along the chromosome and megaplasmid (Fig. 1A). Despite this, all the libraries exhibit two insertion hotspots (Figure S2). One is centered at position 576,160 of the chromosome, disrupting the gene encoding the 5-carboxymethyl-2- hydroxymuconic-semialdehyde dehydrogenase, an enzyme involved in the tyrosine synthesis pathway, and we do not have an explanation for this insertion peak. The second, more frequent hotspot is located at bp 1,457,335 of the chromosome, corresponding to the promoter region of the S-layer gene. This promoter also precedes the kanamycin resistance gene utilized for transposition selection, which may explain this insertion bias. Also, it is worth noticing here that we did find insertions in the Primpol locus and another 13.5 Kb region comprising the AddAB locus. This is was not expected because, as mentioned above, both these regions are deleted in the Δ*ppol/addAB* mutant (genes highlighed with grey background in Table S3), that was the primary host strain to generate the transposition library. A possible explanation for the presence of transposons in these regions is that we used HB27 wild-type strain DNA in the *in vitro* transposition reaction and, upon subsequent transformation of the Δ*ppol/addAB* strain, those regions had the opportunity to get recombinationally repaired with the wild-type DNA at the same time as they were selected for presence of the transposon.

**Figure 1.**
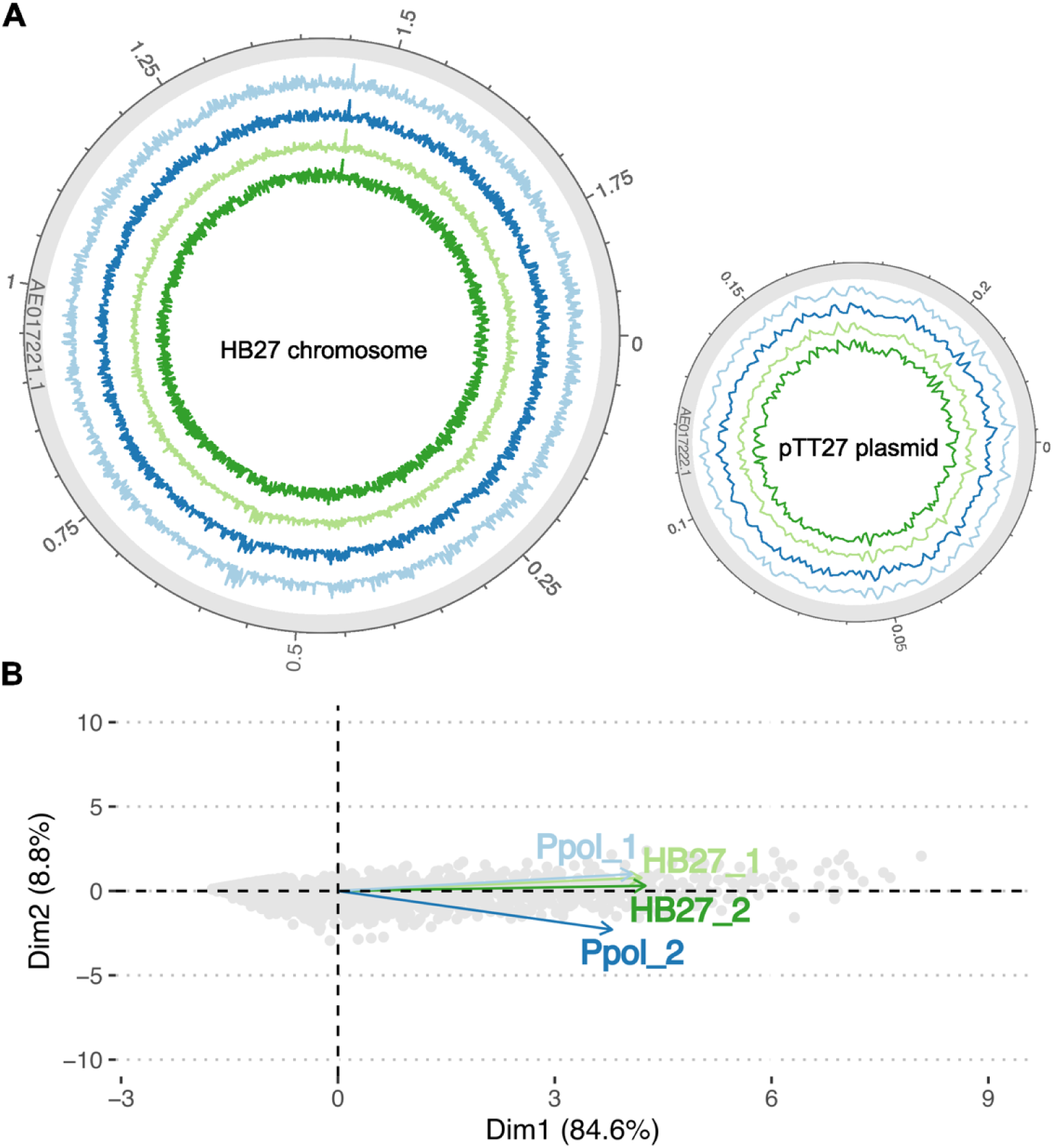
Mapping of Tn-seq Libraries on HB27 genome. **A.** The Tn-seq libraries of HB27 (green and light green) and Ppol (blue and light blue) were mapped onto the HB27 chromosome (left circle) and the pTT27 plasmid (right circle). The lines display the mean coverage per 1 kb windows (in log10 scale). The outer circle includes a genome scale in Mb. **B.** Principal Component Analysis (PCA) of the per-gene Z-score per library. PC1, which accounts for the greatest portion of the variance, primarily reflects the Z-score values, while PC2 explains the sample-to-sample variability. The tight clustering of samples along both principal components suggests a high degree of homogeneity within the dataset.

**Table 1.** Tn-seq insertion sites per library. Only insertions within the 80% central part of the gene were considered.

| Library | Final Reads | Insertions (80% CDS) | Unique Insertions (80% CDS) | Ratio Unique insertions | Normalized. Insertions per M reads |
| --- | --- | --- | --- | --- | --- |
| *Ppol_1* | $9.00 \times 10^5$ | $1.05 \times 10^5$ | $3.69 \times 10^4$ | 35.2 % | $4.10 \times 10^4$ |
| *Ppol_2* | $1.70 \times 10^6$ | $1.64 \times 10^5$ | $4.39 \times 10^4$ | 26.7 % | $2.58 \times 10^4$ |
| *HB27_2* | $1.30 \times 10^6$ | $2.88 \times 10^5$ | $7.14 \times 10^4$ | 24.8 % | $5.49 \times 10^4$ |
| *HB27_1* | $2.70 \times 10^6$ | $1.56 \times 10^5$ | $3.77 \times 10^4$ | 24.1 % | $1.40 \times 10^4$ |
| *Mean* | $1.65 \times 10^6$ | $1.78 \times 10^5$ | $4.75 \times 10^4$ | 27.7 % | $3.39 \times 10^4$ |

The insertions per gene were trimmed eliminating the 10% from both the 5’ and the 3’ end of sequence of the corresponding gene to discard the effect of short N- or C- terminal deletions of the gene product that might not give a clear phenotype. The average number of insertions within the 80% central portion of the gene was around 1.8x10^5^, with roughly 4.7 x10^4^ unique insertion sites (Table 1), resulting in an average of one unique insertion site per 45 bp. The insertion counts per gene were normalized for length of the gene and the total sample reads (see Materials and Methods) and subsequently transformed into log2 scale and centered to a Z- score parameter to facilitate the inter-library comparison. A principal component analysis (PCA) of gene Z-score variability showed high correlation across all samples, with only slight differences between the Ppol replicates (Figure 1B).

### Tn-seq insertion rate does not allow discrimination of essential genes

We evaluated the Z-score of each gene to determine its level of essentiality. When arranging the Z-scores in descending order, we observed a smooth, continuous decline with a downward step at the end, indicating a very small number of genes with very low or no insertions, which would likely correspond to essential genes (Figure 2A). Correspondingly, the density of Z-scores exhibited a unimodal distribution, making it difficult to set a threshold for essentiality (Figure 2B). Various Various previous studies have fit gamma distributions to the bimodal patterns observed in insertion indices (Goodall et al. 2018; Ramsey et al. 2020; Moule et al. 2014; Higgins et al. 2020; Ghomi et al. 2024). However, due to the absence of a clear bimodal curve marking essential genes in our dataset, an unsupervised clustering technique was selected to identify genes with fewer Tn-seq insertions. K- means and PAM clustering methods were compared, as well as the DBSCAN algorithm, which has recently been shown to enhance essentiality classification when bimodal curves are present (A. Ghomi et al. 2024). For our dataset, the K- means method grouping genes into two clusters produced higher average silhouette values, indicating better cluster definition (Figure S3). The resulting clusters were designated as “Highly permissive” (466 genes) and “Intermediate”, corresponding approximately to genes with Z-scores greater than and less than 0.55, respectively. Additionally, we subsequently selected a subset comprising 20% of the Tth genes with a lower insertion rate (453 genes), corresponding to genes with Z-scores approximately below -0.65 across all samples, which we referred to as "Less Permissive". This subset should be considered a subgroup within a range of genes in the “Intermediate” cluster (Figure 3A), useful for data representation and discussion.

**Figure 2.**
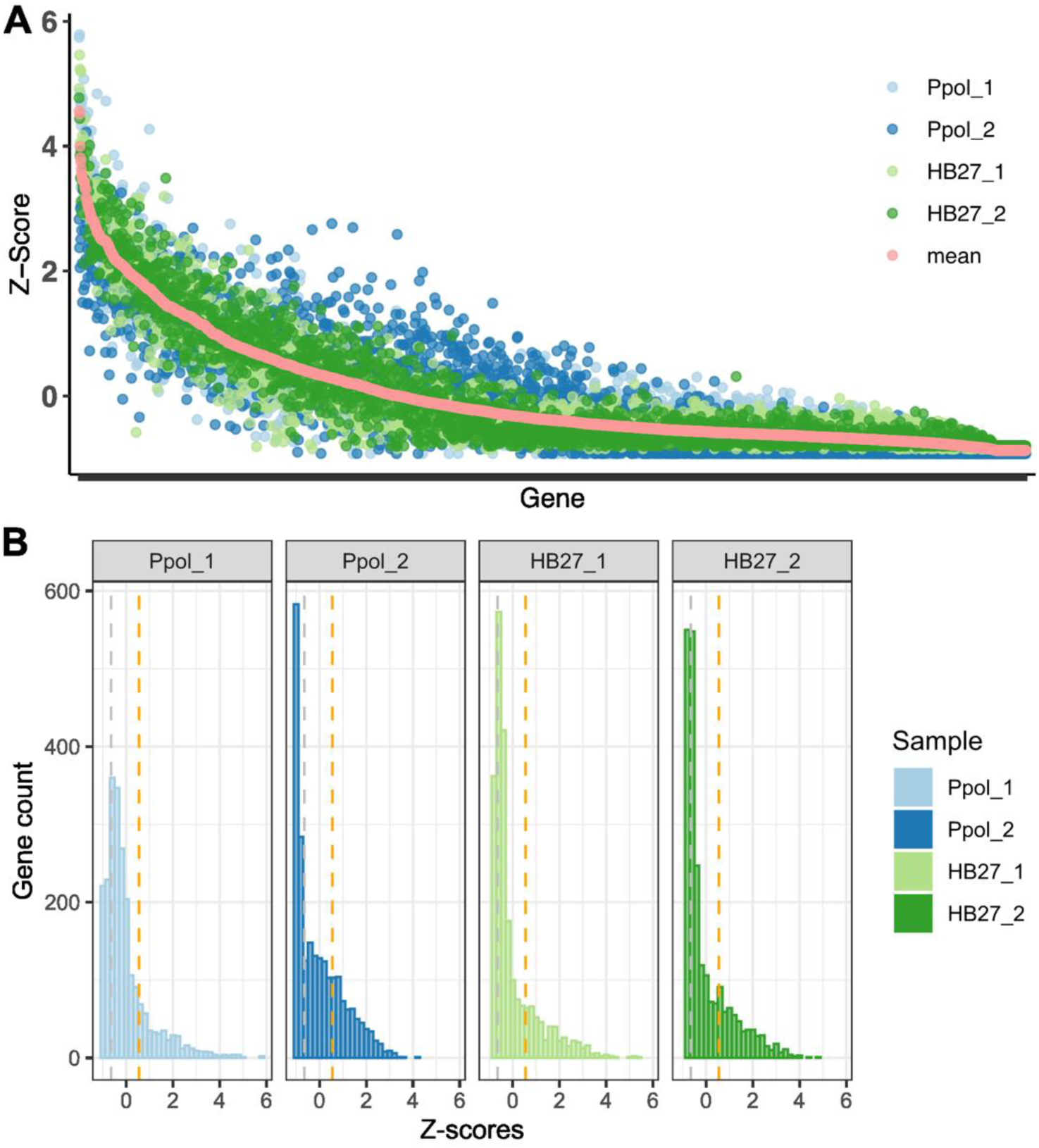
Tn-seq insertion Z-scores per gene. **A.** Z-scores for each gene in each sample, ordered by the average value across all samples (shown in coral). **B.** Distribution of genes based on Z-scores in each library and overall average. Orange and grey dashed vertical lines indicate the Z-scores of 0.55 and - 0.65, which approximately delineate the threshold between Highly Permissive and Intermediate clusters, and the top 20% of genes with fewer insertions across all samples within the permissive cluster (Less Permissive subgroup).

**Figure 3.**
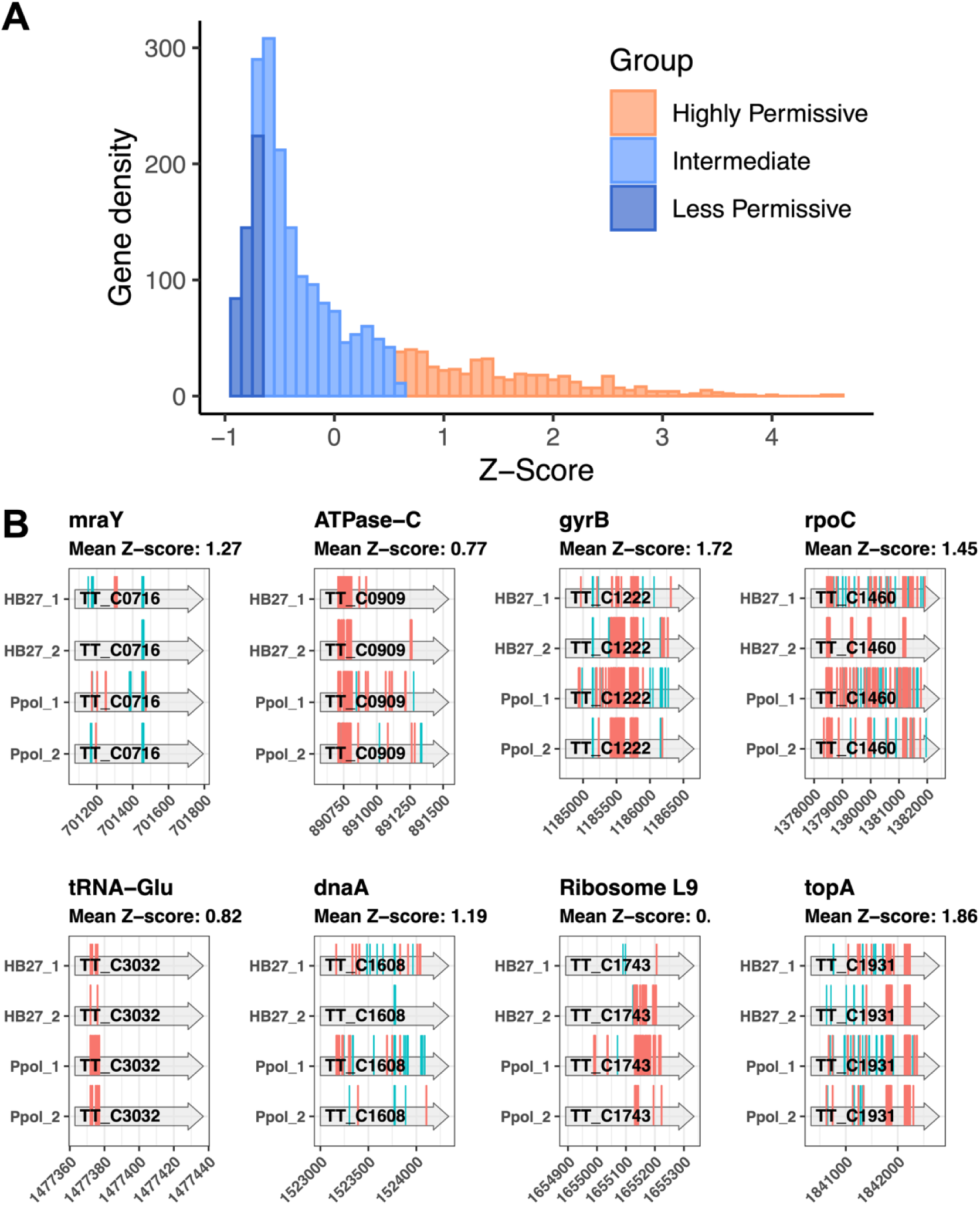
Permissiveness of *Thermus thermophilus* genes to Tn-seq insertions and essentiality. A. Distribution of genes in each Tn-seq group. Note that the “Less Permissive” groups were shortlisted from the “Intermediate” cluster obtained by K-means method. See text for details. B. Detail of detected Tn-seq insertions in selected examples of key biological genes expected to be essential and classified within the “Highly Permissive” group. Insertions in the forward and reverse strand are marked in cyan and magenta, respectively.

The Less Permissive subgroup predominantly comprises genes associated with essential biological functions, such as gene translation and transcription, DNA replication and repair, cell cycle and cell division, metabolism and cell motility (Figure S4A-B), as well as genes of unknown function, being protein synthesis and riboflavin metabolism the only KEGG Orthology functional categories significantly enriched among Less Permissive shortlisted genes (Figure S4C). Notably, several genes expected to be essential were found within both the Intermediate and Highly permissive clusters. These included, for example, those coding for the component of the peptidoglycan synthesis route (*mraY*), the ATP synthase (ATPase-C), a subunit of the DNA gyrase (*gyrB*), a subunit of the RNA polymerase (*rpoC*), non- redundant tRNAs (tRNA-Glu), the replication initiator (*dnaA*), the ribosome subunit (*rpL9*), or the DNA topoisomerase (*topA*), among others (Figure 3B).

### Gene conservation in related taxa does not correlate with Tth Tn-seq results

To put our Tn-seq analysis in context, we constructed pangenomes for *Thermus thermophilus* species, the *Thermus* genus, *Thermaceae* family and Deinococcota phylum using PPanGGOLIN pipeline (Gautreau et al. 2020). This approach enabled us to identify a more sensitive set of conserved or ‘persistent’ gene families compared to recent *Thermus* genus pangenomes (Jiao et al. 2022; Tripathi et al. 2017). As defined in the PPanGGOLIN pipeline, shell genes are those conserved between some individuals of a group, but not most, and cloud genes are rare and found only in one or a few individuals. As expected, the number of persistent gene families increases as we move down in the taxonomic hierarchy (Figure 4A), ranging from 801 to 1744. However, the proportion of Tn-seq Highly Permissive, Intermediate, and Less Permissive HB27 genes remained relatively constant across all taxonomic levels (Figure S5), with approximately 25% of persistent gene families classified as Highly Permissive in all pangenomes (Figure 4B). Thus, the comparison of the three Tn-seq categories with the pangenome analysis results for strain HB27 revealed a low correlation between the Tn-seq groups and pangenome data (Figure 5). This suggests that functional conservation, as inferred from the pangenome analysis, may not be a main determinant of Tn-seq permissiveness.

**Figure 4.**
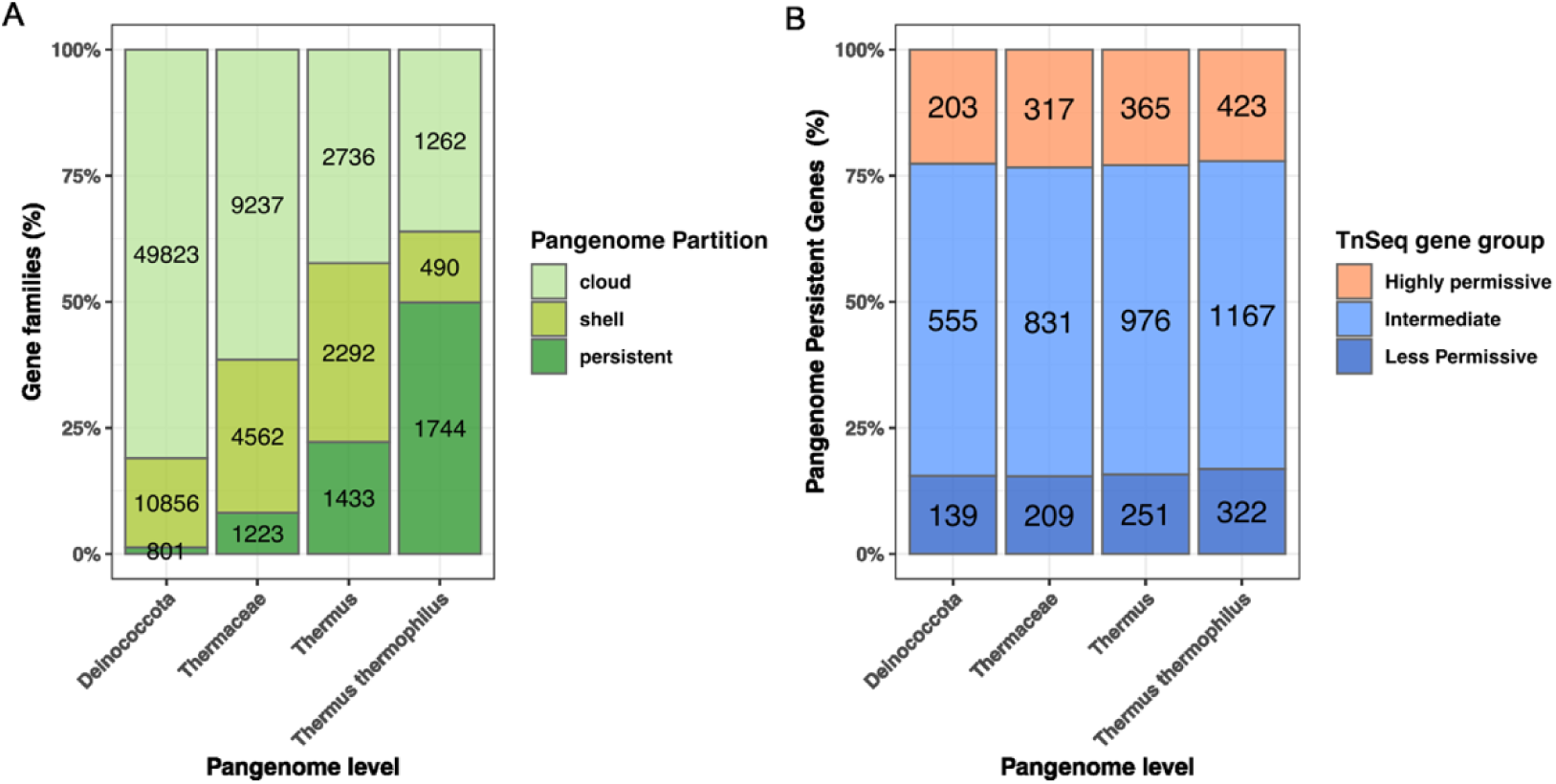
Pangenome contextualization of Tn-seq gene group. A. Pangenome distribution for each taxonomic level, including the number of gene families in each partition is indicated. Pangenomes were obtained with 40% identity and 50% coverage. B. Distribution of persistent (core) gene families from HB27 in each pangenome among the Tn-seq genes groups. See text for details.

**Figure 5.**
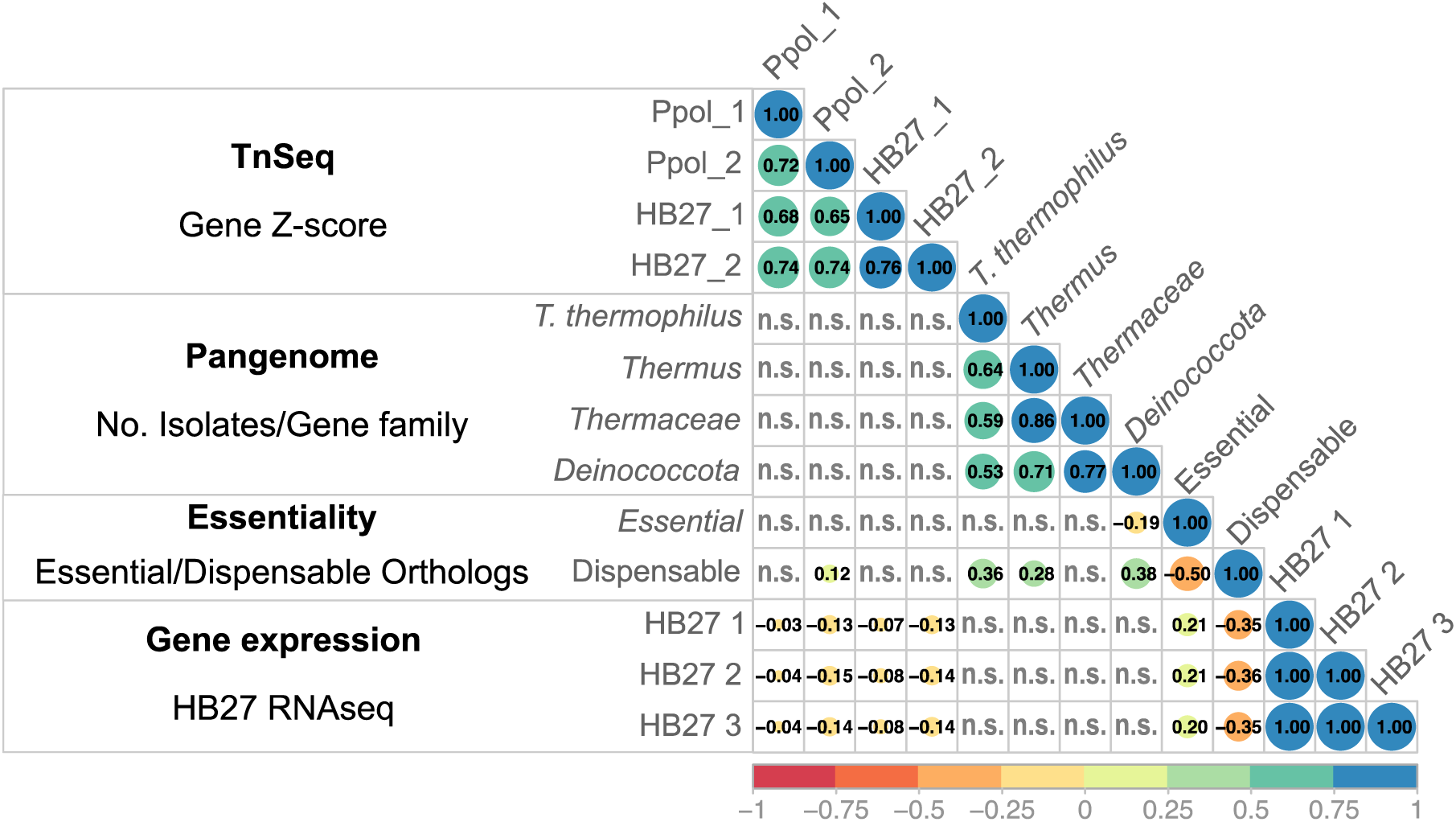
Cross comparison of *T. thermophilus* Tn-seq results. The plot displays the correlation of HB27 Tn-seq Z-scores for each gene with the size (number of isolates) of the corresponding gene family in the pangenomes, the number of conserved ortholog essential/dispensable genes in model bacteria genomes (Rancati et al. 2018), and the HB27 gene expression levels (Swarts et al. 2015). Spearman correlation coefficients for significant correlations (p<0.01) are presented (n.s. stands for non-significant). See Methods section for further details.

Given the lack of clear correlation between gene conservation and the Tn5- insertion tolerance, we analyzed our Tn-seq and pangenomes results in the context of previously published comparative studies on gene essentiality determination. First, we used the comprehensive essentiality data from Rancati et al. (2018), which provides a comparative analysis of the “essentialome” across diverse organisms obtained using different methodologies. We selected the essentiality of *Escherichia coli*, *Mycobacterium tuberculosis*, *Mycoplasma genitalium*, *Pseudomonas aeruginosa*, *Synechococcus elongatus* and *Streptococcus pyogenes* to analyze the correlation between gene essentiality and our Tn-seq results using eggNOG gene ortholog annotations. As shown in Figure 5, essential genes show no correlation with the Tn-seq scores, but exhibit a moderate correlation with gene conservation in pangenomes, which increase at higher taxonomic levels. We obtained similar results when we contextualized our results within the study of Zhang et al. (2018), which includes the essential/non-essential genes of a range of prokaryotic organisms, comprising extremophilic archaea (*Sulfolobus islandicus* and *Methanococcus maripaludis*) and various mesophilic bacteria (*Bacillus subtilis*, *Bacteroides fragilis*, *E. coli*), as well as the minimal essential genome of *Mycoplasma* JVCI Syn3.0. There are 53 genes essential for all these genomes across domains, comprising genes involved mostly in genetic information flux and cell metabolism, which are also frequent functions within Less Permissive genes in our Tn-seq (Figure S4). However, in line with the previous results, about 64% of Tth orthologs of consensus essential genes in the above mentioned genomes were classified as Highly Permissive (6 genes) or Intermediate (27 genes) in our HB27 Tn-seq, but they are, again, highly conserved, as all of them are persistent gene families in the *T. thermophilus* pangenome and near 95% are still members of the core genome of Deinococcota pangenome (Table 2, Table S3).

**Table 2.** Tth TnSeq and Pangenome analysis of consensus prokaryotic essential genes from diverse previous works (Zhang et al. 2018). See Methods for details.

| eggNOG | Description | TnSeq | *T.thermophilus* | *Thermus* | *Thermaceae* | *Deinococcota* |
| --- | --- | --- | --- | --- | --- | --- |
| COG0008 | Glutamate--tRNA ligase | Intermediate | persistent | persistent | persistent | persistent |
| COG0008 | Glutamine--tRNA ligase | Intermediate | persistent | persistent | persistent | persistent |
| COG0013 | Alanine--tRNA ligase | Intermediate | persistent | shell | shell | shell |
| COG0016 | Phenylalanine--tRNA ligase<br>alpha subunit | Highly<br>Permissive | persistent | persistent | persistent | persistent |
| COG0017 | Aspartate--tRNA(Asp/Asn)<br>ligase | Intermediate | persistent | persistent | persistent | persistent |
| COG0017 | Asparagine--tRNA ligase | Less<br>Permissive | persistent | persistent | persistent | persistent |
| COG0018 | Arginine--tRNA ligase | Intermediate | persistent | persistent | persistent | persistent |
| COG0024 | Methionine aminopeptidase | Less<br>Permissive | persistent | persistent | persistent | shell |
| COG0048 | 30S ribosomal protein S12 | Less<br>Permissive | persistent | persistent | persistent | persistent |
| COG0049 | 30S ribosomal protein S7 | Less<br>Permissive | persistent | persistent | persistent | persistent |
| COG0051 | 30S ribosomal protein S10 | Less<br>Permissive | persistent | persistent | persistent | persistent |
| COG0052 | 30S ribosomal protein S2 | Intermediate | persistent | persistent | persistent | persistent |
| COG0060 | Isoleucine--tRNA ligase | Intermediate | persistent | persistent | persistent | persistent |
| COG0081 | 50S ribosomal protein L1 | Intermediate | persistent | persistent | persistent | persistent |
| COG0087 | 50S ribosomal protein L3 | Intermediate | persistent | persistent | persistent | persistent |
| COG0088 | 50S ribosomal protein L4 | Less<br>Permissive | persistent | persistent | persistent | persistent |
| COG0089 | 50S ribosomal protein L23 | Intermediate | persistent | persistent | persistent | persistent |
| COG0090 | 50S ribosomal protein L2 | Less<br>Permissive | persistent | persistent | persistent | persistent |
| COG0091 | 50S ribosomal protein L22 | Less<br>Permissive | persistent | persistent | persistent | persistent |
| COG0094 | 50S ribosomal protein L5 | Less<br>Permissive | persistent | persistent | persistent | persistent |
| COG0096 | 30S ribosomal protein S8 | Less<br>Permissive | persistent | persistent | persistent | persistent |
| COG0097 | 50S ribosomal protein L6 | Intermediate | persistent | persistent | persistent | persistent |
| COG0098 | 30S ribosomal protein S5 | Less<br>Permissive | persistent | persistent | persistent | persistent |
| COG0099 | 30S ribosomal protein S13 | Less<br>Permissive | persistent | persistent | persistent | persistent |
| COG0100 | 30S ribosomal protein S11 | Less<br>Permissive | persistent | persistent | persistent | persistent |
| COG0102 | 50S ribosomal protein L13 | Intermediate | persistent | persistent | persistent | persistent |
| COG0103 | 30S ribosomal protein S9 | Less<br>Permissive | persistent | persistent | persistent | persistent |
| COG0124 | Histidine--tRNA ligase | Less<br>Permissive | persistent | shell | persistent | persistent |
| COG0162 | Tyrosine--tRNA ligase | Highly<br>Permissive | persistent | persistent | persistent | persistent |
| COG0172 | Serine--tRNA ligase | Less<br>Permissive | persistent | persistent | persistent | persistent |
| COG0180 | Tryptophan--tRNA ligase | Intermediate | persistent | persistent | persistent | persistent |
| COG0186 | 30S ribosomal protein S17 | Less<br>Permissive | persistent | persistent | persistent | persistent |
| COG0195 | Transcription<br>termination/antitermination<br>protein NusA | Intermediate | persistent | persistent | persistent | persistent |
| COG0197 | 50S ribosomal protein L16 | Less<br>Permissive | persistent | persistent | persistent | persistent |
| COG0198 | 50S ribosomal protein L24 | Intermediate | persistent | persistent | persistent | persistent |
| COG0199 | 30S ribosomal protein S14 type Z | Less Permissive | persistent | persistent | persistent | persistent |
| COG0201 | Protein translocase subunit SecY | Intermediate | persistent | persistent | persistent | persistent |
| COG0202 | DNA-directed RNA polymerase subunit alpha | Intermediate | persistent | persistent | persistent | persistent |
| COG0256 | 50S ribosomal protein L18 | Intermediate | persistent | persistent | persistent | persistent |
| COG0361 | Translation initiation factor IF-1 | Intermediate | persistent | persistent | persistent | persistent |
| COG0442 | Proline--tRNA ligase | Highly Permissive | persistent | persistent | persistent | persistent |
| COG0462 | Ribose-phosphate pyrophosphokinase | Highly Permissive | persistent | persistent | persistent | persistent |
| COG0462 | Ribose-phosphate pyrophosphokinase | Intermediate | persistent | persistent | persistent | persistent |
| COG0470 | DNA polymerase III subunit delta' | Intermediate | persistent | persistent | persistent | shell |
| COG0470 | Belongs to the ribF family | Intermediate | persistent | shell | shell | shell |
| COG0480 | Elongation factor G | Highly Permissive | persistent | persistent | persistent | persistent |
| COG0480 | Elongation factor G | Intermediate | persistent | persistent | persistent | persistent |
| COG0495 | Leucine--tRNA ligase | Highly Permissive | persistent | persistent | persistent | persistent |
| COG0504 | CTP synthase | Intermediate | persistent | persistent | persistent | persistent |
| COG0522 | 30S ribosomal protein S4 | Less Permissive | persistent | persistent | persistent | persistent |
| COG0525 | Valine--tRNA ligase | Intermediate | persistent | persistent | persistent | persistent |
| COG0532 | Translation initiation factor IF-2 | Intermediate | persistent | persistent | persistent | persistent |
| COG0592 | Beta sliding clamp | Intermediate | persistent | persistent | persistent | persistent |

Finally, correlation analysis with available HB27 gene expression data (Swarts et al. 2015) showed a low correlation between expression levels and ortholog gene essentiality (r^2^ values of 0.2 and -0.36 for essential and dispensable genes, respectively, from Rancati et al. 2018), but a slight negative correlation with the Tn-seq Z-score. Thus, highly transcribed genes in HB27 are enriched within the highly conserved (persistent) group and, overall, they would get somewhat less hits in our Tn-seq. Altogether, these results indicate that despite the different lifestyles and significant phylogenetic distances among the genomes examined in prior essentiality studies, the expected higher conservation of universally essential genes is applicable to Tth, even though, in this case, the Tn-seq approach may not readily disclose most of essential genes.

### Essential genetic pathways accumulate Tn5 insertions

Some of the single copy genes conserved across prokaryotic genomes, operating separate functions of the cell, and known to be generally essential are being clustered as Highly permissive or Intermediate in our Tn-seq (Figure 3B and Table S3). While, at the same time, the other components of the corresponding routes fall into the Intermediate or Less Permissive categories, which is what would be expected for important cellular machineries such as the ribosome, RNA polymerase or DNA replisome (Figure S4). To exemplify the fact that, even within complexes that would be expected to be essential, as a whole, for Tth, genes classified as Less Permissive are intermixed with genes labeled as Highly Permissive by Tn-seq, we show the scheme for respiratory Complex I (Figure 6). This multisubunit complex should have a stoichiometric composition and, as the major entry point for electrons from NADH into the electron transport chain, should also be essential or very important for the strictly aerobic Tth. Despite its critical role, we observe that nearly two-thirds of its subunits are encoded by Intermediate or Highly Permissive genes, with only a reduced portion of Less Permissive components.

**Figure 6.**
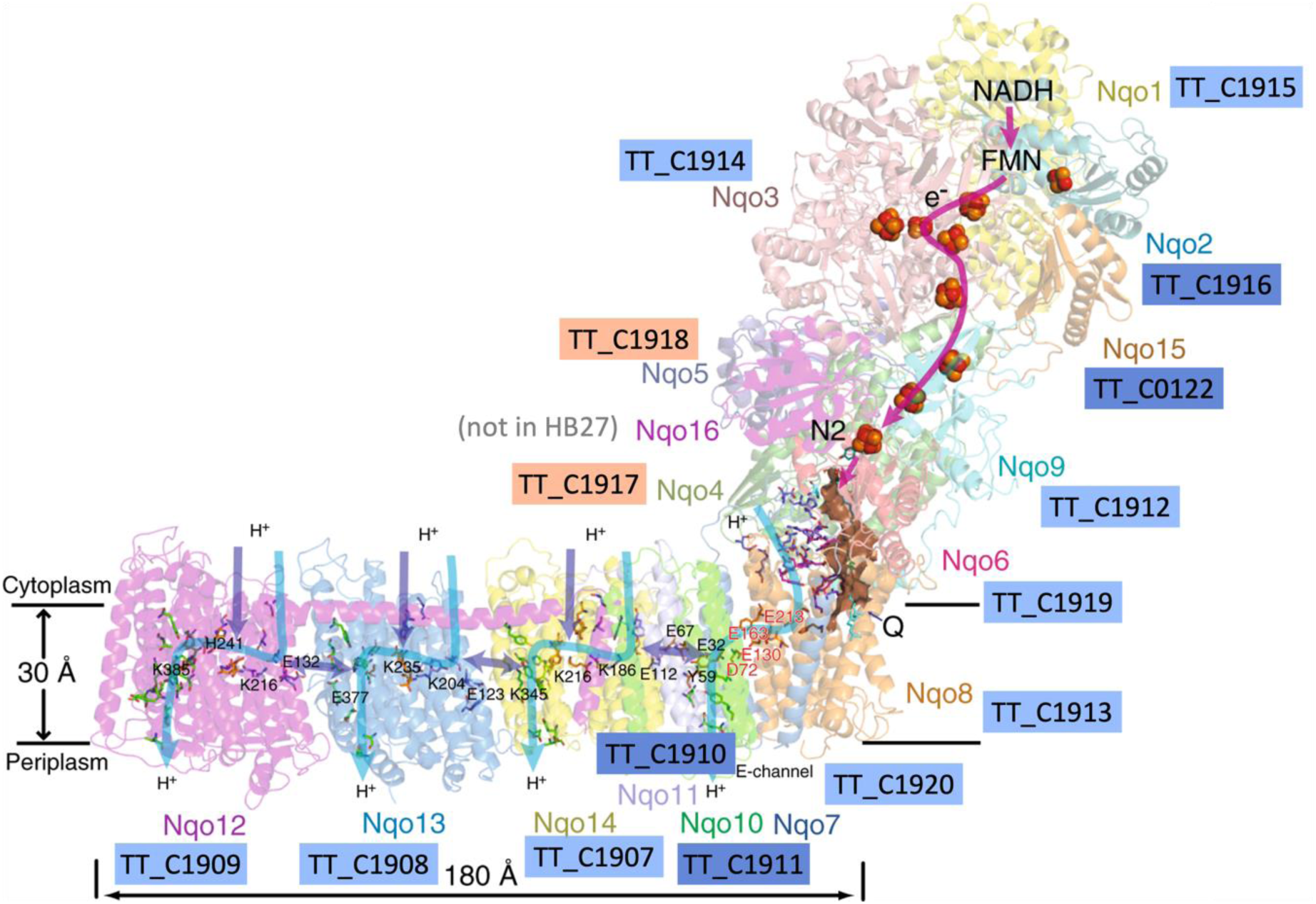
Respiratory Complex I proteins are encoded by many genes within the Intermediate or Highly Permissive Tn-seq groups. Representation of the crystallographic structure of respiratory Complex I (encoded by *nqo* genes) of *Thermus thermophilus* HB8 (Berrisford, Baradaran, and Sazanov 2016). The equivalent genes in Tth HB27 are indicated with the color code that correspond with Less Permissive (cobalt), Intermediate (sky blue) or Highly Permissive (peach) Tn-seq groups.

Similarly, although genes related with ribosomal protein synthesis are enriched in the Less Permissive group, as expected for essential functions, more than 40% of rRNA and ribosomal protein genes are classified as Intermediate or even Highly Permissive, as it is the case for *rpsR* and *rplI*, which encode the small subunit proteins L9 and S18, respectively (Table S3).

### Redundancy of DNA repair routes is compatible with Tn-seq insertion permissiveness

As mentioned in introduction DNA repair routes are expected to be important and conserved in a microorganism living at extreme temperatures such as Tth. In Figure 7, we have summarized the indicated DNA repair routes, the Tn-seq data, the pangenome classification and the available experimental data on gene dispensability under standard, non-stress conditions. Interestingly, the overall impression is that most of the genes are, at some degree, individually dispensable. This conclusion is based on experimental results, and supported by the fact that the majority of these genes fall in the Intermediate or Highly Permissive groups, with the only exception of one dispensable gene (*nfo*) (Nakane et al. 2012), which is classified as Less Permissive according to the Tn-seq analysis. On the other hand, genes involved in several routes and generally considered essential, such as those coding for the ligase LigA or RecA, are labelled as Intermediate by Tn-seq. Notwithstanding, a *recA* deletion mutant has been reported for Tth (Castan et al. 2003), the resultant strain, however, is genetically very unstable, so, for this analysis, we consider *recA* a quasi-essential gene. Curiously, the Tth SSB gene falls into the Intermediate category even when there is only one paralog/copy and is generally considered part of the bacterial minimal core genome (Glass et al. 2006). Other genes encoding proteins involved in several repair pathways, such as RadA, RecN, RarA, DdrA, DprA or PcrA, are classified as Intermediate or Highly Permisive, which is consistent with existing knowledge on these genes in other organisms. The *addAB* and *ppol* genes Tn-seq data is affected by the fact that these genes are deleted from the Ppol starting strain; therefore, we have excluded them from the Tn-seq classification. However, previous work from our laboratory has shown that these genes appear to be dispensable only when certain compensatory mutations occur simultaneously in the genome (Verdú et al. 2022). Most DNA repair genes are highly conserved and belong to the persistent pangenome categories, with the exception of *nfo*, *udg*, *udgB*, *yfjP*, *recQ* and *pcrA*. Interestingly, *ppol* and *addAB* are conserved in *Thermus* but absent from *Deinococcus radiodurans* (*D.r.*). We also observed that in the recombinational repair section, four helicase-nuclease pairs appear to be present: *addAB*, *recJ/recQ*, *nurA/hepA* and *sbcCD* that have been shown to be individually dispensable (Shimada et al. 2010; Fujii et al. 2018; Blesa et al. 2017), with the exception of *sbcCD*, for which no deletion data are available. Similarly, there are at least two resolvase-type pathways, RuvABC and RecG, with *ruvB* known to be dispensable. In the RecFOR pathway, on the other hand, *recR* is dispensable, *recO* is conditionally dispensable, and no *recF* deletion mutant could be obtained (Gómez-Campo et al. 2024), and all of them cluster as Intermediate by Tn-seq. The *recQ* and *pcrA* genes, located on the pTT27 plasmid, which is known to be almost entirely dispensable (Naoto Ohtani, Tomita, and Itaya 2016), are classified as Highly Permissive and Intermediate, respectively, and they belong to the shell cluster in all the pangenomes.

**Figure 7.**
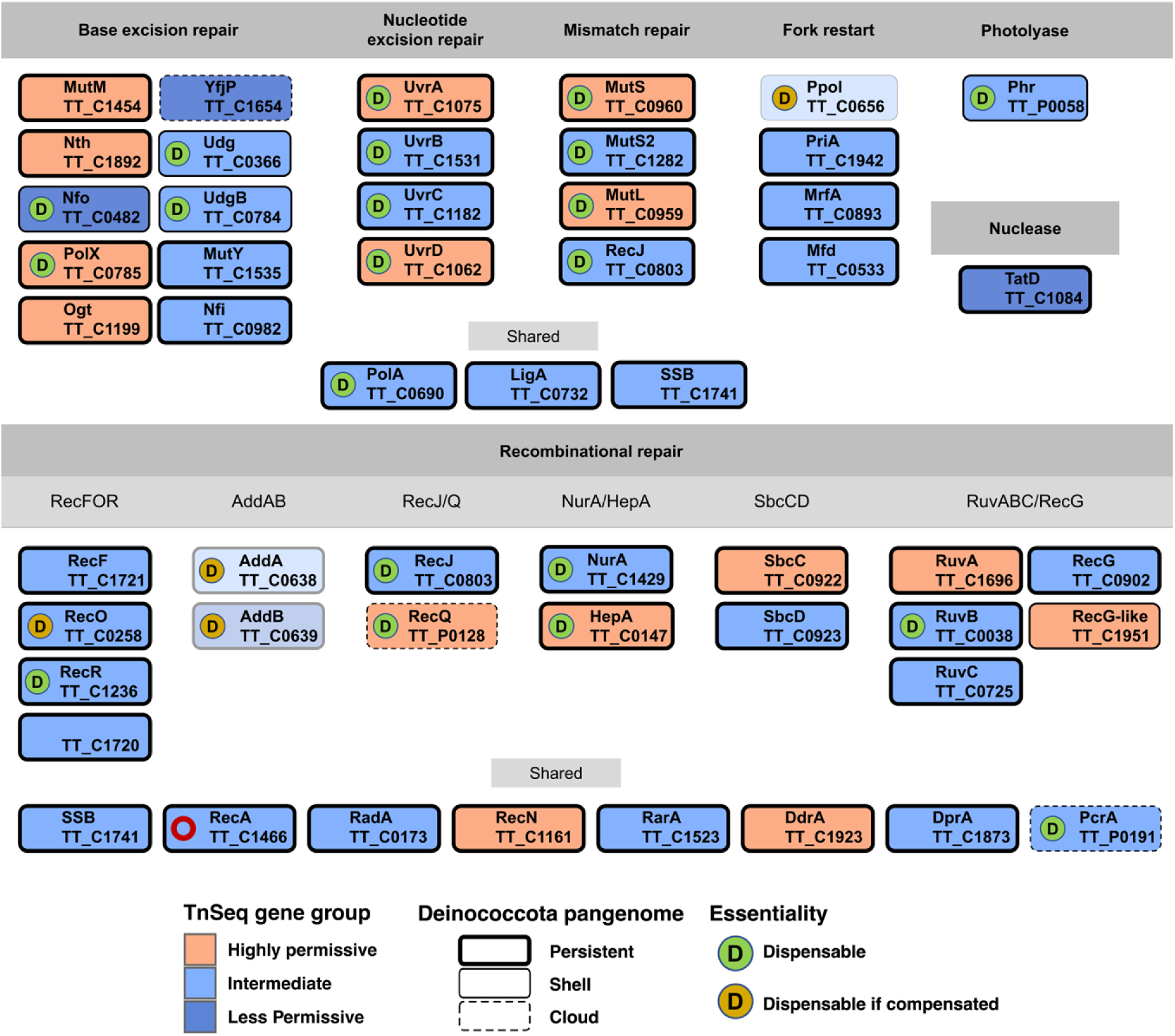
DNA repair proteins encoded by *Thermus thermophilus* HB27. The proteins are grouped according to repair route. The gene names are shown below the protein names (previous nomenclature). As indicated, the classification derived from Tn-seq data follows the color code: Highly Permissive (peach), Intermediate (sky blue) or Less Permissive (cobalt) Tn-seq groups. The Deinococcota pangenome classification is denoted by the thickness of the corresponding rectangle, from persistent (thick line), to shell (thin line) to cloud (dashed line). Dispensable genes (under non-stress conditions), based on available experimental evidence, are denoted with a D in a green circle or in an orange circle if the deletion mutants have compensatory mutations. The red circle in RecA denotes that the deletion mutant is very unstable. AddA/B and Ppol genes are shown semitransparent because they were absent in the Δppol recipient strain.

Altogether, the Tn-Seq insertions patterns in DNA repair genes and the existing experimental data suggest that Tth does not rely heavily on a single repair pathway, but more probably on the simultaneous activity of several routes. Thus, at least in the case of recombinational repair, there appears to be redundancy of pathways, as it observed in other bacteria. Interestingly, for the relatively small genome of Tth, the panoply of encoded repair functions suggests a preference for recombinational, error-free DNA repair, as exemplified by the presence of several helicase-nuclease pairs. A comparison with the *D. r.* repair mechanisms seems to support that idea since, for example, Tth lacks canonical translesion synthesis DNA polymerases, which are present in *D.r.* (Blanchard and De Groot 2021).

### Hypothetical proteins and DUFs

All genomes have a proportion of genes encoding proteins without assigned function, although many of them contain conserved domains, referred to as Domains of Unknown Function (DUFs). In our Tn-seq analysis, roughly 20% of proteins with unknown functions belong to the Less Permissive category. We performed Alphafold DB and DALI model-based structural predictions to attempt the assignment of putative functions to the selected domains. The models predicted at least general functions for most DUFs (Table S4), however, in most cases, it remained unclear why those DUFs with their corresponding functions would be important for the cell to the level that they had low transposon hit rate.

For example, the DUF1931 (TT_C1115)-encoded predicted protein showed strong structural homology with histone folds, however these prokaryotic domains lack the conserved histone DNA-binding residues. Recently, structural approaches confirmed the existence of these histone variants encoded by DUF1931-containing genes, suggesting that this domain in prokaryotes may have another possible functions beyond genome organization (Schwab et al. 2024).

In any case, we have assembled a catalog of putative functions for DUFs, which remains open for further research to more precisely determine the biological roles of these domains.

## Discussion

The size of bacterial genomes ranges from those containing a few hundred genes to more than eleven thousand. As more information is accumulated on the ecology, physiology, metabolism and gene expression in bacteria, the gene complement of sequenced strains can be assigned to express the proteins that perform the functions or capabilities attributed to the corresponding organism. As mentioned in introduction, it has been shown that there is a minimal set of genes, present in the immense majority of bacteria, even in strict intracellular parasites, that represent the core functions required for bacteria to survive. Starting from this minimal set, the next ampliation would be defined by what is required for a free-living organism considering its metabolism. Beyond that point, additional sets of functions or capabilities imply a corresponding increase in the amount of encoding DNA. *Thermus thermophilus* is a free-living bacterium with a streamlined genome that harbors the capabilities to live at high temperature. Our Tn-seq study adds an extremophilic and polyploid bacterium to the few present in the list of genomes analyzed using this technique, and represents a new tool that can be used for different purposes.

The method we used is based on the fact that this strain is naturally competent and recombination proficient. However, to increase the initial diversity of the library, we used a mutant with greatly enhanced capacity of transformation efficiency and DNA fragment incorporation to its genome. In addition, we isothermally amplified the product of the Tn5 transposition reaction to further increase the number of recombinants obtained. With this, an adequate number of initial transformants was achieved, and the same strategy could likely be applied to other bacteria as long as they can be made competent and support recombination with sufficient efficiency.

After processing the deep sequencing data, we expected to observe a bimodal distribution in which one of the peaks would correspond to near-zero insertions (i.e., essential genes), and the other would be a distribution from moderate to a high number of insertions (i.e., very permissive genes). However, in our case, all the samples exhibited unimodal distributions that resembled the second peak of the bimodal distribution. Thus, there are no genes without insertions, and we used unsupervised clustering to group the genes, obtaining two main groups, a wide range group of Highly Permissive genes and a group of Intermediate genes, which we operatively split to obtain a subset of Less Permissive genes and Intermediate genes. When we compared the Less Permissive group of quasi-essential genes (453 genes) with the core of conserved genes across bacterial species, we found a series of coincidences in some of the expected essential functions (Fig. S4), but surprisingly there was a substantial number of key genes classified as Intermediate or even Highly Permissive, such as those encoding the SSB, DnaA, several ribosomal proteins, tRNAs, subunits of the RNA polymerase, peptidoglycan synthesis genes, ATPase, and others. In no other study have been observed so many conserved genes to be Highly Permissive to transposon insertions. A possible explanation for the permissiveness of genes typically considered essential is that Tth has been reported to be polyploid, with 5 to 8 copies of the genome at all times. With this, one can imagine a scenario in which, for a given gene, the transposon is present only in a fraction of the genome copies within a cell. If the gene is essential, the transposition would never colonize the rest of the genomes, reaching an equilibrium between the number of resistance cassettes and the copies of the essential gene required for viability/growth. Thus, in each cell division, the daughter cells, should randomly inherit at least one copy of the resistance cassette and the essential gene to remain viable, and then return to the equilibrium situation. When performing a gene deletion in Tth by recombinative insertion of a resistance cassette, we routinely manage to completely eliminate the target gene when it is not essential, simply by successive re-streaking the transformants on selective plates. However, when the gene is essential, we observe in some cases small transformant colonies that, upon streaking on selective plates, this is, forcing the expansion of the cassette, fail to continue growing. This situation ressembles what we observe by Tn-seq, where individual, isolated colonies, with insertion in important genes may exhibit some apparent growth, and therefore that gene would show as allowing some degree of permissivity after the Tn-seq procedure, yet the mutant would not be viable in a sustained way. We have only found one example of Tn-seq essentiality analysis in polyploid bacteria, the cyanobacteria *Synechococcus elongatus* PCC 7942, which detected a pool of essential genes in the first peak of a bimodal density curve (Rubin et al. 2015). However, it is actually an oligoploid organism with 3-4 genome copies per cell (Griese, Lange, and Soppa 2011), which would reduce the plasticity and tolerance to Tn-seq insertions. Interestingly, a 6000-clone initial Tn-5 transposition library was reported for *D. r.*, also a polyploid bacterium (Hansen 1978). After selection for sensitivity to genotoxic agents, 208 insertion mutants were studied (Dulermo et al. 2015), of which half were heterozygous, and the authors showed that some of the heterozygous mutants had insertions in essential genes such as *dnaE*, *dnaN*, *holA*, *topA* and *gyrB*, which lends support to polyploidy as a key factor in interpreting our results.

Continuing with our case, after insertion of a Tn5 cassette within an important gene in a cell, the number of copies of the encoded protein present in the cell would depend on the stability of the RNA, the stability of the protein, the number of genomes with the insertion, the regulation of the expression, and other factors, all this in a dynamic fashion. As the cells divide, the daughter cells will have to regain equilibrium. One can imagine that a protein that is itself very stable, depends on a very stable mRNA, and/or has a self-regulatory circuit, would tolerate being in fewer genomic copies. As a result, it potentially could have more copies with the inserted cassette and, thus, appear as Intermediate or even Highly Permissive after Tn-seq. This could be the case for DnaA or SSB, with high protein amounts per cell and with self-regulation. On the contrary, a protein with low stability, low mRNA stability or lacking individual regulation would require more gene copies and therefore, it would appear as Less Permissive. This explanation could also be extended to proteins that form multicomplexes, in the sense that, individually, those proteins may have lower stability or higher replacement rates.

On the other hand, there are genes in our study that appeared as Less Permissive, for which we do not find a functional argument for the lack of permissiveness. We do not have a clear explanation for this, other than suggesting that the insertion of the cassette could have non-predicted negative effects on the expression of genes in the vicinity.

Tth, considered a typical bacterium, still has aspects of its gene regulation that are yet not well-understood. For example, no known mechanism for SOS regulation has been reported for Tth, and it also has several defense systems that could potentially affect gene expression. These or other causes could lead to the type of effects that we are observing, and additional research will be required to shed light on the actual mechanisms involved.

The panoply of DNA repair genes and their functions are a key aspect of Tth biology, that probably reflects its adaptation to a thermophilic lifestyle. As there is a substantial number of genes in this category for which knock-outs have been reported, we have compared the published data on essentiality with the Tn-seq results. Not counting the *ppol* and *addAB* genes that are not present in the Tn-seq starting strain (*ppol/addAB*), out of 23 genes reported as dispensable under non- stressed, rich medium conditions, 22 were labeled as Highly Permissive or Intermediate in our study. Only the dispensable gene *nfo* was labelled as Less Permissive by Tn-seq. Some genes have been shown to be dispensable only when compensatory mutations occur simultaneously in the genome, such as as *ppol* itself and *recO*. In general, it is somewhat curious that many repair genes are non- essential in Tth, considering that the high temperature probably induces higher levels of DNA damage, which would require adequate repair in real time. As mentioned above, and as it happens in other bacteria, redundancy in repair factors and routes could probably explain the dispensability of individual genes in this category. The assessment of the repair genes supports the view that our analysis, in general, captures the expected pattern for most genes. However, our study did not yield a set of genes specifically essential in the way other studies do, and we also observed a substantial number of exceptions, in both directions, to what we would predict to be essential or not in this organism.

In conclusion, this new Tn-seq study in *T. thermophilus* introduces a general method to facilitate such studies and provides overall consistent results with the expected functional patterns of gene essentiality, while unexpectedly allowing for multiple exceptions in conserved genes with orthologs among essential genes in diverse bacteria. These findings offer a novel perspective on the assumed correlation between gene disruption tolerance and gene requirement in bacterial genomes, especially in polyploid species, and could potentially be expanded in the future with increased knowledge and study of more microorganisms.

## Experimental Procedures

### Strains and growth conditions

Tth was cultivated in Thermus broth (TB) culture medium (Tryptone 8 g/l, Yeast extract 4 g/l, NaCl 3 g/l, and NaOH 1.2 ml/l in Sierra de Cazorla brand mineral water at 65°C. Liquid cultures were incubated under aeration (180 rpm) in 50 ml flasks with a final volume of 10 ml. Cultivation in solid TB medium was carried out on TB medium supplemented with 2% (w/v) agar plates in a humidification chamber. Kanamycin (Kn, 30 μg/ml) was used for selection of the mutant library.

The Tth strains used were HB27, as a reference wild-type, and HB27 Δ*ppol* (Verdú et al. 2022), as a high transformation efficiency host.

To obtain a transposable kanamycin resistance cassette the oligonucleotides TnKanPslp and TnKanEnd (Table S1) having at the ends the Mosaic End sequences optimally recognized by the Tn5 transposase (Ez-Tn5 Transposase (Lucigen)) were used to amplify the thermostable resistance encoding gene from plasmid pMK184 (de Grado et al., 1999). This fragment was cloned into plasmid pUC19 between the EcoRI and HindIII sites to obtain plasmid pUCTnKat. This plasmid was used as a template to perform PCR with the oligonucleotides M13 Forward (-20) and M13 Reverse (-27). The product obtained was treated with the DpnI enzyme for 1 hour at 37 °C to eliminate the template plasmid, purified using the “DNA Clean and Concentrator-5” system (Zymo Research) and used directly in the transposition reactions.

In the transposition reaction, 1 unit of the EZ-Tn5 transposase with its corresponding reaction buffer, 300 ng of *T. thermophilus* HB27 genomic DNA (prepared according to Marmur (1961)) and 100 ng of the PCR fragment corresponding to the transposon were used in a final volume of 10 µl. The mixture was incubated at 37°C overnight and finally the reaction was stopped by heating at 70°C for 10 minutes. Given that the direct transformation into Tth of the transposition reaction gave rise to a low number of colonies, it was decided to amplify that product isothermally and representatively, using the “Repli-G” kit from Qiagen. The amplification was carried out according to the manufacturer’s instructions using 2.5 µl of the transposition reaction as a template, and the reaction was incubated for 16 hours at 30°C. The reaction was stopped by heating to 65°C and the DNA obtained was fragmented by 10 passes through a 0.5 mm hypodermic needle (21G). 5 µl of the reaction, with a DNA concentration of 18 ng/µl, was used directly in the transformation of 0.5 mL of a culture of *T. thermophilus* Δppol strain at OD600= 0.4. Ten transformations were plated on TB agar kanamycin plates to obtain the Ppol_1 initial library of 144000 transformants. Transformation of Tth was performed according to Koyama et al., (1986). The plates were incubated at 60 °C in a humid chamber for approximately 48 hours and then colonies were pooled by suspension in TB medium and pools were frozen and kept at -20 °C for further use. To obtain genomic DNA aliquotes of 2 ml of cells at an OD600 of 3 were used for DNA extraction with the NZYTech NZY Microbial gDNA Isolation kit. For subsequent transformation to obtain libraries HB27_1, HB27_2 or Ppol_2, 100 ng of genomic DNA from Ppol_1 extraction were used in transformations as described above.

Tn-seq sequencing was carried out at the Genomics facility of the Madrid Science Park (https://fpcm.es/genomica/#servicios), using the MiSeq system and the Illumina MiSeq Reagent Kit v3 for 150-cycle single-end sequencing.

To generate the library, a first fragmentation of the genome was done, followed by ligation of the Index Fork adapter (Table S1) at their ends. To amplify the sequence associated with the transposon insertion, a first PCR was performed with the oligonucleotide KatH, directed to the 3’ end of the *kat* gene sequence, and with Index Primer R, as reverse primer, which by joining only part of the Index Fork adapter sequence, allows amplification to be in a single direction. During the second PCR, the region comprised between the 3’-end ME and the forked sequence was amplified. For this, variants of the P5-InvRep-Var primer were used, composed of the P5 sequence (binding adapter to the sequencing platform), Illumina sequencing primer 1 (Rd1 Sp) and a variable spacer region (spacer) at the 5’ end, followed by the sequence complementary to the ME. P7-AD014-index-R was used as the primer at the 3’ end, with the forked sequence followed by the barcode and the binding adapter to the P7 sequencing platform at the 5’ end of the primer. The oligonucleotides utilized are summarized in Table S1.

### Tn-seq libraries sequencing and gene insertion scoring

A custom pipeline was designed to process the raw sequencing reads and analyze the data to calculate gene insertion scores. Data management, statistical analysis and data plotting were performed with R (v. 4.4.1) in the R Studio suite (v. 2024.12.0). Main specific R packages from CRAN or Bioconductor are indicated. See the available repositories for full details.

Briefly, quality check, reads filtering, trimming and deduplication was carried out with FastP 0.23.4 (Chen et al. 2018) and reports were merged with MultiQC 1.22.3. Reads processing parameters (-q 20 -r --cut_right_window_size 10 -- cut_right_mean_quality 30 -w 10) were optimized to maximal proportion of mapped reads against the reference HB27 assembly GCA_000008125 using Bowtie2 2.5.1 (with --very-sensitive option). Reads mapping breadth and depth was calculated with Bedtools 2.30.0, WeeSam 1.6 and BAMdash 0.3.1.

We considered only Tn-seq insertions derived from reads mapping within the 10- 90% interval of each gene to avoid issues with truncated or chimeric proteins. Tn insertion counts are derived from read mapping coordinates, normalized by the total mapped reads in coding regions. We use a log2 transformation to manage pseudocounts and scale the data with the R function scale() to obtain a Z-score, facilitating sample comparison. Thus, our score will be obtained with the following formula:

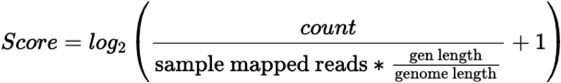

Then, we calculated final Z-score using the R *scale* function:

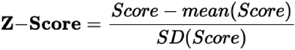

Sample Z-score comparison was carried out by PCA analysis with R package *factoextra.* Plots were obtained with R packages *ggplot, ggenes,* and *ggenomes.* Some of the final plots were modified with Inkscape to improve layout or clarity.

### Tn-seq gene classification and correlation with pangenome

In order to classify the genes by the ratio of Tn5 insertions, we compared PAM, Kmeans and DBSCAN clustering methods using the R packages *cluster*, *factoextra* and *dbscan*, respectively. The best results by highest average silhouette width were obtained for a Kmeans clustering in two main groups (Figure S3). Then, the large group with less insertion rate was split into two groups by shortlisting the 20% of genes with lower Z-score, obtaining the three groups of Less Permissive, Intermediate and Highly Permissive genes. Gene functions as well as COG and KEGG functional categories were then annotated with EggNog mapper (Cantalapiedra et al. 2021). Functional enrichment was assessed with R package *ClusterProfiler*.

To generate pangenomes at species, genus, family and phyla taxonomic levels, we used NCBI datasets tool (v.16.5.0) to download the available RefSeq genomes (May 27th, 2024). After uniform reannotation with Bakta (v. 1.9.2, using full database v. 5.1), pangenomes were constructed with PPanGGOLIN (v. 2.0.5) using MMSeqs clustering sequence identity and coverage parameters to 0.4 and 0.5. For integration of RefSeq HB27 annotation (GCA_000008125.1) with the fresh Bakta annotation, we used a flexible table merge with *difference_inner_join* function in R *fuzzyjoin* package, allowing the maximal distance for joining that did not give rise to duplicate features (*max_dist*=210).

For comparison of Tn-seq data with available data of essential genes and HB27 transcriptome analysis, datasets from Zhang et al. (2018), Roncati et al. (2018) and Swarts et al. (2015), respectively, were downloaded and parsed for uniform format and comparison with custom R scripts available in the GitHub repository. Correlation analysis was carried out with package *corrplot*.

## Data availability

Raw sequencing libraries and code for bioinformatic analyses have been deposited in e-CienciaDatos repository under DOI: doi.org/10.21950/AJSMCI

A detailed description and code of Tn-seq analysis is also available online in a GitHub repository (https://github.com/mredrejo/thermus_tnseq) and a Quarto report in https://mredrejo.github.io/thermus_tnseq/.

## Acknowledgements

This work was supported by grants from the Spanish Ministry of Science, Innovation and Universities [TED2021-130430B-C22] and [PID2022-137468OB-I00] to BS, JB, and MM. CG and MG had a contract from Fundación Severo Ochoa. An institutional grant from Fundación Ramón Areces to the CBMSO is also acknowledged. MRR lab was funded by MCIN/AEI/10.13039/501100011033 and ERDF A way of making Europe [PID2021-123403NB-I00].

